# Progression and Resolution of SARS-CoV-2 Infection in Golden Syrian Hamsters

**DOI:** 10.1101/2021.06.25.449918

**Authors:** Kathleen R. Mulka, Sarah E. Beck, Clarisse V. Solis, Andrew L. Johanson, Suzanne E. Queen, Megan E. McCarron, Morgan R. Richardson, Ruifeng Zhou, Paula Marinho, Anne Jedlicka, Selena Guerrero-Martin, Erin N. Shirk, Alicia Braxton, Jacqueline Brockhurst, Patrick S. Creisher, Santosh Dhakal, Cory F. Brayton, Rebecca T. Veenhuis, Kelly A. Metcalf Pate, Petros C. Karakousis, Sabra L. Klein, Sanjay K. Jain, Patrick M. Tarwater, Andrew S. Pekosz, Jason S. Villano, Joseph L. Mankowski, for the Johns Hopkins COVID-19 Hamster Study Group

## Abstract

To catalyze SARS-CoV-2 research including development of novel interventive and preventive strategies, we characterized progression of disease in depth in a robust COVID-19 animal model. In this model, male and female golden Syrian hamsters were inoculated intranasally with SARS-CoV-2 USA-WA1/2020. Groups of inoculated and mock-inoculated uninfected control animals were euthanized at day 2, 4, 7, 14, and 28 days post-inoculation to track multiple clinical, pathology, virology, and immunology outcomes. SARS-CoV-2-inoculated animals consistently lost body weight during the first week of infection, had higher lung weights at terminal timepoints, and developed lung consolidation per histopathology and quantitative image analysis measurements. High levels of infectious virus and viral RNA were reliably present in the respiratory tract at days 2 and 4 post-inoculation, corresponding with widespread necrosis and inflammation. At day 7, when infectious virus was rare, interstitial and alveolar macrophage infiltrates and marked reparative epithelial responses (type II hyperplasia) dominated in the lung. These lesions resolved over time, with only residual epithelial repair evident by day 28 post-inoculation. The use of quantitative approaches to measure cellular and morphologic alterations in the lung provides valuable outcome measures for developing therapeutic and preventive interventions for COVID-19 using the hamster COVID-19 model.

## Introduction

In December 2019, a novel beta coronavirus was isolated from patients that presented with severe and ultimately fatal pneumonia in Wuhan, China^1^. The virus was designated Severe Acute Respiratory Syndrome Coronavirus 2 (SARS-CoV-2) and rapidly spread through human-to-human transmission, causing the current global pandemic of Coronavirus Disease 2019 (COVID-19). As of June 2021, there have been over 168 million confirmed cases and over 3.5 million deaths globally attributed to SARS-CoV-2 infection^2^.

While many organ systems can be affected by SARS-CoV-2 infection, pulmonary disease has been most frequently associated with severe and fatal cases of COVID-19^3^. The earliest stage of disease is characterized by edema and vascular damage including endothelial cell degeneration and necrosis, with neutrophilic infiltration of alveolar septa and capillaries (endothelialitis and capillaritis) and microthrombosis^3–6^. This is followed by an exudative phase of diffuse alveolar damage (DAD), with fibrinous edema in the alveolar spaces, increased numbers of macrophages and epithelial multinucleated giant cells, hyaline membrane formation, and epithelial necrosis followed by type 2 pneumocyte hyperplasia. Additionally, vascular changes occur, including endothelial necrosis, hemorrhage, thrombosis of capillaries and small arteries, and vasculitis^5,7^. In turn, the organizing stage of DAD, and the final fibrotic stage of DAD ensue which may include proliferation of myofibroblasts within the lung interstitium, and deposition of collagen leading to fibrosis. Squamous metaplasia has also been observed^3,8^.

The emergent and widespread nature of this pandemic necessitated the rapid development of multiple animal models and biologic systems to study various aspects of pathogenesis, treatment, and prevention of disease. To date, reported animal models of COVID-19 pathology include human ACE2 transgenic mice^9–12^, golden Syrian hamsters^12–18^, nonhuman primates^19,20^, and ferrets^21,22^; recent comprehensive reviews of animal models of COVID-19 were provided by Zeiss et al., and Veenhuis and Zeiss, 2021^23,24^. Each model species has advantages and limitations with respect to similarity to disease in humans, expense, and practicality. The hamster model offers several advantages over other animal models: it is a relatively small, immunocompetent animal that is susceptible to infection with varied SARS-CoV-2 clinical isolates and readily develops pulmonary disease. Specifically, hamsters consistently develop moderate to severe bronchointerstitial pneumonia characterized by acute inflammation, edema, and necrosis 2-4 days post SARS-CoV2 challenge, progressing to proliferative interstitial pneumonia with type II pneumocyte hyperplasia by 7 days post-challenge. Pulmonary lesions have been reported to resolve around 10-14 days post-inoculation with little to no evidence of residual damage ^13–18,20,25^.

Although several studies have provided an overview of pulmonary pathology during acute infection, comprehensive longitudinal assessments of pulmonary pathology are lacking, including chronic time points. Likewise, there is a dearth of information available integrating clinical, pathology, virology, and immunology findings or reporting systemic pathologic findings associated with SARS-CoV-2 infection in hamsters. In this study, we provide in-depth, longitudinal, pathological characterization of multisystemic disease manifestation caused by SARS-CoV-2 infection in male and female golden Syrian hamsters. Furthermore, we objectively measured tissue damage and inflammatory responses by digital image analysis using an open-source platform, QuPath^26,27^. Our results show that inoculating hamsters intranasally with SARS-CoV-2 reliably induces acute damage to the respiratory tract with initial viral replication followed by a macrophage-dominant pulmonary immune response. In turn, a reparative phase follows with abundant type II pneumocyte hyperplasia restoring the alveolar lining, mirroring SARS-CoV-2 infection in humans.

## Materials and Methods

### Animals

7 to 8 week old male and female golden Syrian hamsters *Mesocricetus auratus* (Envigo) were singly housed in the JHU CRB Animal Biosafety Level 3 (ABSL)-3 facility. After acclimation (7 day minimum), hamsters were sedated intramuscularly with xylazine and ketamine then inoculated intranasally with 10^5^ TCID_50_ of SARS-CoV-2 USA-WA1/2020, (BEI Resources NR#52281) diluted in 100uL DMEM; virus stocks. Uninfected animals were mock-inoculated with 100 μL of DMEM intranasally to serve as controls. Groups of 12 animals (4 mock and 8 SARS-CoV-2 inoculated, equal numbers of males and females) were euthanized at 2, 4, 7 and 14 days post inoculation (DPI). An additional group of 4 mock (2 male and 2 female) and 19 SARS-CoV-2-inoculated hamsters (10 male and 9 female) were euthanized at 28 DPI. Body weights were measured daily until 10 DPI, then on 14, 21, and 28 DPI. Blood samples were collected at 0, 7, 14, 21, and 28 DPI depending on group. Terminal blood samples obtained via cardiac puncture were saved for serology, coagulation assays, FACS analysis, hematology, and clinical chemistries. At study endpoints, animals were euthanized with intraperitoneal sodium pentobarbital. A complete post-mortem examination with comprehensive tissue harvest (flash frozen and 10% NBF immersion fixed samples) was performed on all hamsters. ACUC statement. The animal procedures in this study were in accordance with the principles set forth by the Institutional Animal Care and Use Committee at Johns Hopkins University and the National Research Council’s Guide for the Care and Use of Laboratory Animals (8^th^ edition)

### SARS-CoV-2 measurements

Infectious virus titers in the respiratory tissue homogenates were determined by the 50% tissue culture infectious dose (TCID_50_) assay^28^. Briefly, tissue homogenates were serially diluted 10-fold, transferred in sextuplicate into 96-well plates confluent with Vero-E6-TMPRSS2 cells (obtained from the Japan Institute of Infectious Diseases), incubated at 37^0^C for 4 days, and then stained with naphthol blue-black solution for visualization. Infectious viral titers (TCID_50_/mL) were determined by Reed and Muench method. For detection of SARS-CoV-2 genome copies, RNA was extracted from lungs using the Qiagen viral RNA extraction kit (Qiagen) and reverse transcriptase PCR (RTqPCR) was performed as previously described^29^. Viral data from a subset of these animals has been included in a report on humoral responses in this model^28^.

### Postmortem examination

Standard necropsy procedures included gross examination of all major organs. After obtaining organ weights, samples were either frozen or immersion fixed in 10% neutral-buffered formalin. The lungs were weighed en bloc, and then the left lobe was removed at the level of the left bronchus. Formalin was gently infused into the left bronchus to inflate the left lobe, then the entire lobe was submerged in formalin. The right lung lobes were divided into different frozen sample collection tubes for preparing tissue homogenates and RNA extraction.

### Histopathology analyses

After immersion fixation in 10% neutral-buffered formalin for 72 hours, tissues were trimmed, processed, and embedded in paraffin. 5µm thick sections were mounted on glass slides and stained with hematoxylin and eosin (H&E). Histopathological analysis was performed independently by two veterinary pathologists (S.B. and K.M.) blinded to animal identification and infection status. Tissues evaluated included nasal cavity, trachea, lung (left lobe), esophagus, stomach, small intestine, cecum, large intestine, brain, heart, kidney, liver, gallbladder, spleen, adrenal gland, reproductive organs, urinary bladder, lymph nodes, salivary glands, bone, haired skin, skeletal muscle, bone marrow, and decalcified cross sections of the head.

### *In situ* hybridization and immunohistochemistry

#### *In situ* hybridization to detect SARS-CoV-2 RNA

*In situ* hybridization (ISH) was performed on 5µm-thick sections of formalin-fixed lung mounted on charged glass slides using the Leica Bond RX automated system (Leica Biosystems, Richmond, IL). Heat-induced epitope retrieval was conducted by heating slides to 95°C for 15 minutes in EDTA-based ER2 buffer (Leica Biosystems, Richmond, IL). The SARS-CoV-2 probe (cat. 848568, Advanced Cell Diagnostics, Newark, CA) was used with the Leica RNAScope 2.5 LS Assay-RED kit and a hematoxylin counterstain (Leica Biosystems, Richmond, IL). Slides were treated in protease (Advanced Cell Diagnostics, Newark, CA) for 15 minutes and probes hybridized to RNA for 1 minute. An RNApol2 probe served as a hamster gene control to ensure ISH sensitivity; a probe for the bacterial dap2 gene was used as a negative control ISH probe.

#### Immunohistochemistry

10% NBF-fixed lung sections of SARS-CoV-2 infected and control animals were immunostained with anti-Iba-1 antibody (1:2000; Wako; 019-19741 Richmond, VA), anti-CD3 antibody (1:200; DAKO; Ref: A0452 Santa Clara, CA, or anti-pan-cytokeratin antibody (1:1000; Santa Cruz; sc-8018 Dallas, TX). Heat-induced epitope retrieval was conducted by heating slides to 95°C for 20 minutes in sodium citrate-based ER1 buffer (Leica Biosystems, Richmond, IL) before immunostaining. Dual immunostaining for the epithelial marker pan-cytokeratin and the macrophage marker Iba-1 was performed on lung sections of SARS-CoV-2-infected animals. For epitope retrieval, slides were heated to 100°C in sodium citrate-based ER1 buffer for 20 minutes (Leica Biosystems, Richmond, IL). Slides were then stained with anti-Iba1 antibody (1:2000; Wako; 019- 19741 Richmond, VA) using the Bond Polymer Refine Red Kit (cat. DS9390 Leica Biosystem, Richmond, IL). The slides were stained using a pan-cytokeratin antibody (1:1000; Santa Cruz; Sc-8018; Dallas, TX) with the Bond Polymer Refine Kit (cat. DS9800 Leica Biosystem, Richmond, IL). Immunostaining was performed using the Bond RX automated system (Leica Biosystems, Richmond, IL). Positive immunostaining was visualized using DAB and RED, and slides were counterstained with hematoxylin.

### Digital Image Analysis

Whole slides containing sections of the entire left lung lobe cut through the long axis were scanned at 20x magnification on the Zeiss Axio Scan.Z1 platform using automatic tissue detection with manual verification. Lung sections were analyzed using QuPath v. 0.2.2 (complete code included in supplemental methods). To measure Iba-1 and CD3 immunostaining, the create thresholder function was applied to detect levels of DAB above a threshold that was designated as positive within a given annotated area, or region of interest (ROI). The percent positive ROI was calculated using positive area quantitated by the thresholder divided by total area of the ROI. For SARS-CoV-2 ISH quantitation, the train pixel classifier tool was used. Within an ROI, annotations were created and designated as either positive or ignore, which allowed QuPath to correctly identify areas of positive staining. Percent positive ROI was calculated using positive area detected by the classifier divided by total area of the ROI.

To quantitate consolidation of 7 DPI lungs, the wand tool was used to outline each scanned section of lung, creating an annotation. Superpixels were generated using the DoG superpixel segmentation function in QuPath. These detections were then selected, and intensity features were added. Next, areas within the ROI were annotated using multiple slides of infected and control animals that were designated as “Consolidation”, “Non-consolidated”, “Atelectasis”, or “Ignore”. This allowed the classifier to successfully detect areas of affected tissue, while ignoring areas that were unaffected, densely stained due to normal tissue architecture, or collapsed due to variable formalin infusion of the lungs. Percent consolidation was calculated using the number of superpixels identified as consolidated divided by the total number of superpixels detected for a given slide.

### Flow Cytometry

Freshly collected whole blood in EDTA was stained for FACS analysis with the limited antibodies available that cross-react in the hamster model at the initiation of the study. Cell surface markers for CD4 (anti-mouse, clone GK1.5, Biolegend, San Diego, CA), CD8 (anti-rat, clone 341, BD biosciences, San Jose, CA), MHC II (anti-mouse, clone 14- 4-4S, BD Biosciences), and a Live/Dead discriminator (Fixable Near-IR Dead Cell Stain Kit, ThermoFisher, Waltham, MA) consistently stained populations of lymphocytes or monocytes allowing for a limited understanding of the proportion of these cell types. Whole-blood samples were stained with pre-titered amounts of the indicated monoclonal antibodies using 50ul of whole blood at room temperature for 20 min. Whole blood samples were then lysed and fixed in 2 ml of FACS Lysing Solution (BD Biosciences, San Jose, CA) for 10 min at room temperature. Samples were collected in a centrifuge at 400 x g for 5min, washed in 2 ml of 1x phosphate-buffered saline (PBS), and then resuspended in 0.5 ml of PBS for analysis. Flow cytometry was performed on a BD LSRFortessa (BD Biosciences, San Jose, CA). Data were analyzed using FlowJo 10.0.8 software (FlowJo, LLC, Ashland, OR). Whole blood was first gated on size and complexity using FSC-A and SSC-A to remove debris, followed by using FSC-H and FSC-A and SSC-H and SSC-A measurements to remove doublet populations. Single cells were then gated into lymphocyte or monocyte populations based on size and complexity profiles, and then gated to include only live cells using a Live/Dead viability stain. Live lymphocyte-sized cells were gated as MHC II negative/dim (T cells) or MHC II+ (B cells). The MHC II-negative cells were further gated on their expression of CD4 or CD8. Populations of MHC II+ B cells, CD4+ lymphocytes, and CD8+ lymphocytes were calculated as a proportion of total viable lymphocytes. Live monocyte-sized cells were further gated as MHC II+.

### Statistical analysis

Statistical analyses were performed using GraphPad Prism version 7.04. One-way analysis of variance (ANOVA) was used to detect significant differences between groups of animals sampled at different time points post-inoculation. The two-sample t-test was used to evaluate differences between two groups. Infectious virus titers and viral RNA copies were log transformed and compared using two-way ANOVA with mixed-effects analysis followed by Bonferroni’s multiple comparison test.

## Results

### Clinical Outcomes

The most robust finding following SARS-CoV-2 inoculation was decreased body weight (**Fig 1A**). Animals inoculated with SARS-CoV-2 progressively lost body weight in the first week of infection until the 6 DPI nadir (−13.8% decline in group mean body weight from baseline; 19.3 % lower than control animal group mean at 6 DPI, P < 0.001) before gradually rebounding over the course of the following 3 weeks. At day 28 DPI, control and infected group body weights were not significantly different (P=0.095). Sex differences in body mass loss have been documented in hamsters infected with SARS-CoV-2, and those differences were explored previously^28^.

**Figure 1.**
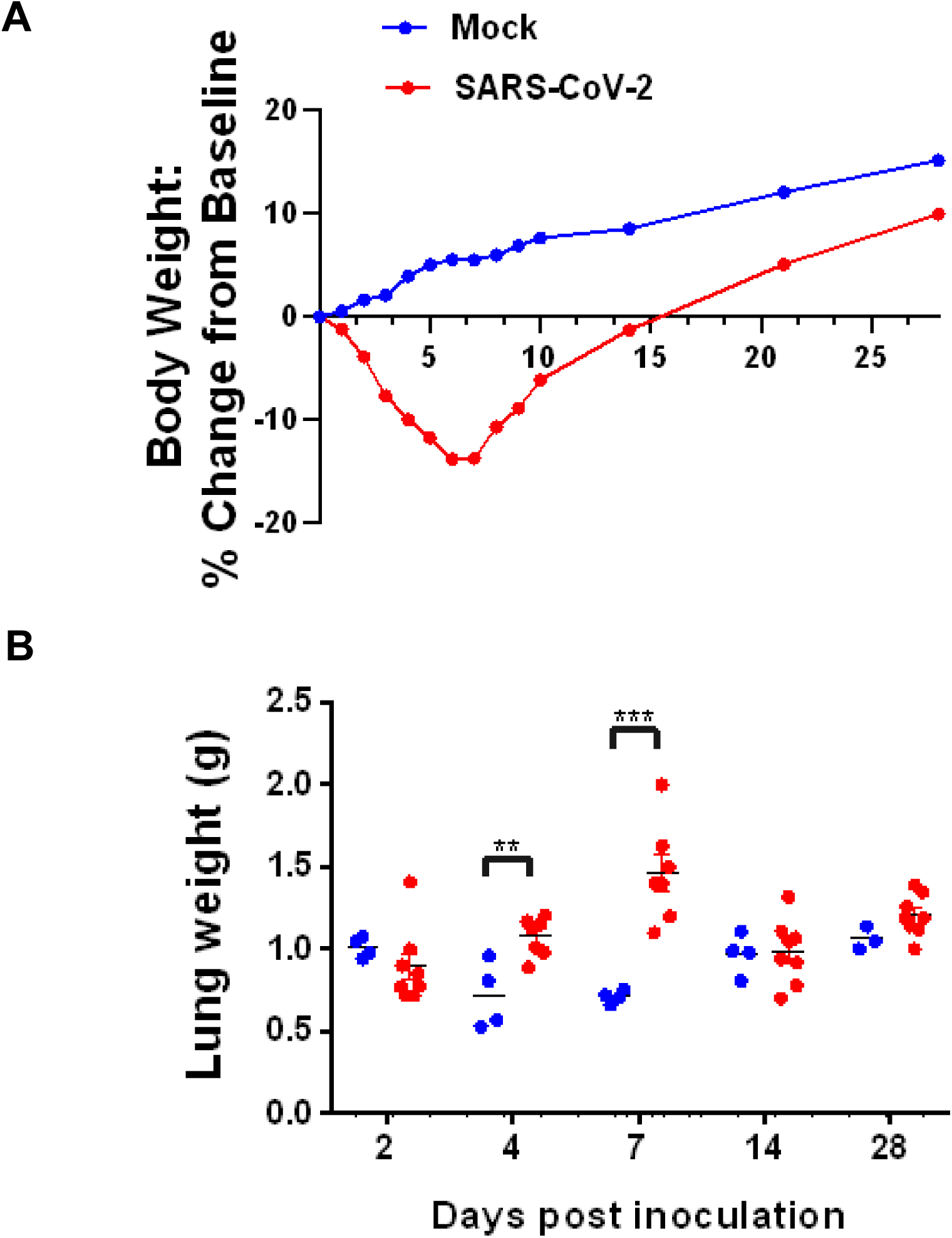
Body weight change over time and postmortem lung weights. A. Body weight of control mock-inoculated hamsters (shown in blue) compared with SARS-CoV-2 inoculated hamsters (red). Mock-inoculated animals demonstrated a steady increase typical of young growing animals. In contrast, SARS-CoV-2 inoculated animals had a sharp decline in body weight until 6 DPI when body weight began to increase. B. At time of necropsy, lung weights of SARS-CoV-2-inoculated (red) hamsters had significantly higher total lung weights at both 4 and 7 DPI (P= 0.004, P= 0.0008, respectively, unpaired t-test) as compared to mock-inoculated animals, consistent with lesion severity and consolidation.

SARS-CoV-2-inoculated hamsters did not consistently develop clinical signs of respiratory disease; only mild nasal discharge or slight increased respiratory effort was observed intermittently in very few animals.

### SARS-CoV-2 infectious virus and viral RNA measurements in the respiratory tract

Peak infectious viral titers in the nasal turbinates, trachea, and lungs were present at 2 DPI, then decreased at 4 DPI (Fig. 2). The highest levels of infectious SARS-CoV-2 were found in nasal turbinates at 2 DPI (5.0 x 10^7^ TCID_50/mL_); lung levels were highest at 2 DPI (7.33 x 10^6^ TCID_50_). While infectious virus was cleared from the respiratory tract of most of the hamsters by 7 DPI, infectious virus was detected in the lungs of a single animal. No sex differences in viral load were detected in infected hamsters^28^.

**Figure 2.**
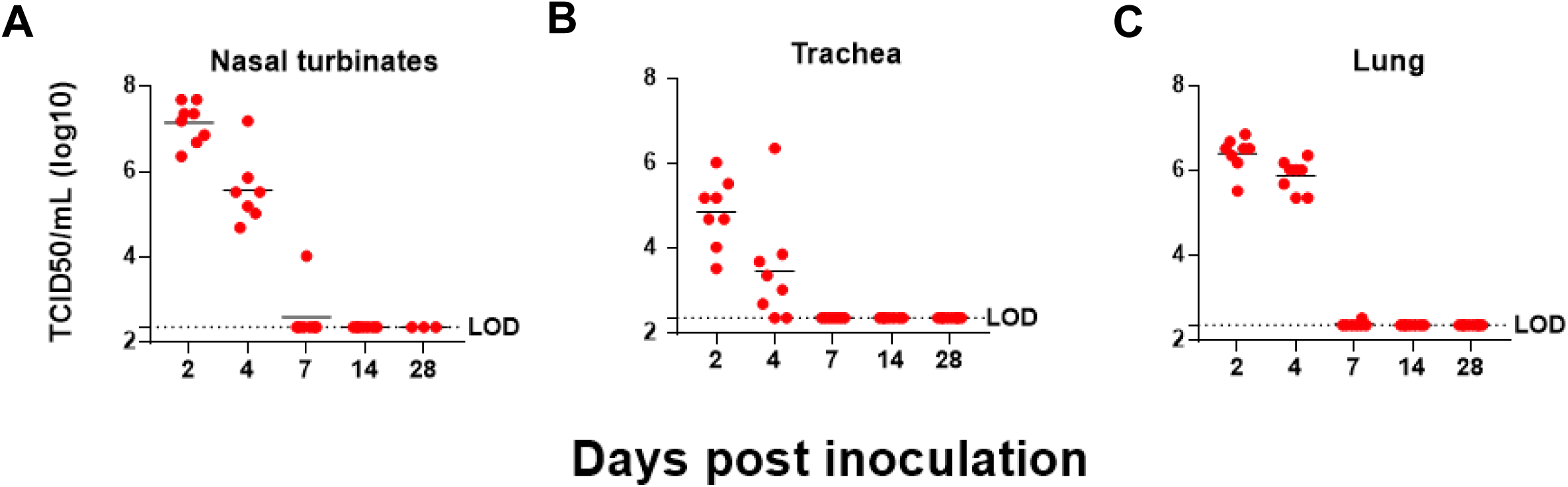
Viral loads throughout the respiratory system. Tissue culture infectious dose 50 (TCID50) assay revealed the highest levels of infectious virus present in the nasal turbinates (A), trachea (B), and lungs (C) at 2 DPI. Detectable levels of infectious ARS- CoV-2 were still present in all these tissues at 4 DPI; only a single animal had low level infectious virus at 7 DPI.

### Gross Pathology

On gross necropsy examination, lungs from both infected and control animals were indistinguishable, with variably mottled red to brown coloration randomly distributed in all lung lobes, suggesting that this change was due to perimortem procedures and not a reliable indicator of pathology related to SARS-CoV-2 infection. Two animals had adhesions between the lung lobes and the diaphragm; both were infected animals euthanized 7 DPI. Compared to uninfected control animals, average post-mortem lung weights were 50% higher in infected animals at 4 DPI and 105% higher at 7 DPI (**Fig 1B**; P = 0.014; P <0.0001, respectively, 2-way ANOVA multiple comparisons). All other tissues were unremarkable in both control and SARS-CoV-2-infected animals.

### Longitudinal Progression of Lesions in the Respiratory System

#### 2 DPI

The nasal cavity contained eosinophilic proteinaceous exudate with abundant degenerate neutrophils in all infected animals (8/8) and no uninfected animals (0/4). In 7 of 8 infected animals, olfactory epithelium damage ranged from degeneration to necrosis with erosion accompanied by infiltrating neutrophils classified as rhinitis. The tracheal submucosa contained mild infiltrates of mononuclear cells in 3/6 infected animals with tissue available for evaluation, (tissue not present in 2/8 infected animals), and no uninfected animals. In infected animals, within the lungs, (Fig 3, 2 DPI) the following changes were observed: minimal to moderate suppurative bronchitis and bronchiolitis (8/8), bronchial and bronchiolar intraluminal neutrophilic necrotic cellular debris (7/8), bronchial epithelial syncytia (2/8), alveolar septa expanded by inflammatory cells and eosinophilic proteinaceous material (4/8), intra-alveolar macrophages, neutrophils, necrotic cellular debris, and fibrinous exudate (6/8). Vascular changes in lungs included: reactive vascular endothelium within small to medium arteries (4/8), perivascular edema, and leukocytes infiltrating through the vascular walls (2/8). The lungs of the uninfected control animals had no significant findings. Intra-alveolar hemorrhage was observed intermittently in both uninfected and infected groups at all time points attributed to tissue collection artifact.

**Figure 3.**
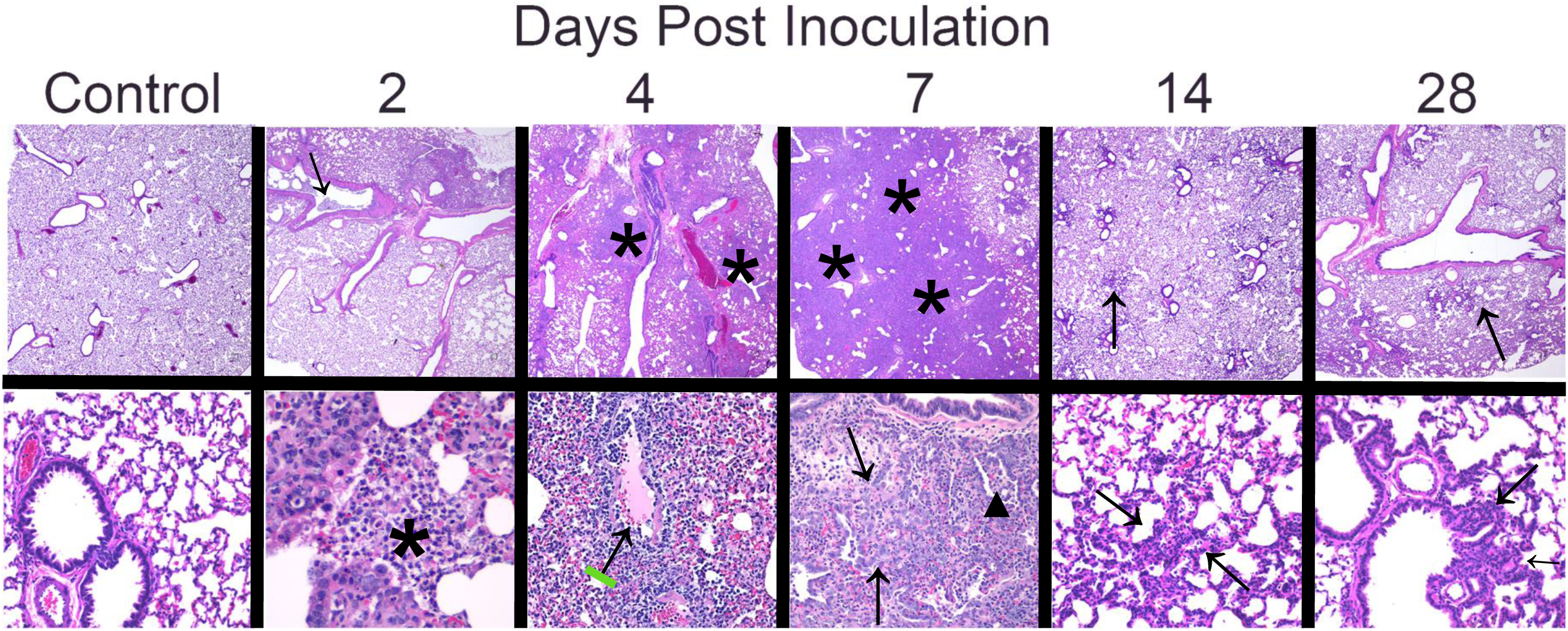
Progressive morphologic alterations in lung over time. Mock-inoculated (intranasal saline alone) animals that served as uninfected controls had no lesions identified by histologic examination. 2DPI: Lung lesions included intraluminal neutrophilic infiltrates and necrotic cellular debris (black arrow), as well as abundant intra-alveolar macrophages, neutrophils, necrotic cellular debris (black asterisk), and fibrinous exudate. 4DPI: Lesions included areas of consolidation consisting of type II pneumocyte hyperplasia, numerous intra-alveolar macrophages, neutrophils, necrotic cellular debris, and eosinophilic fibrinous exudate (black asterisk), and multifocal vasculitis with reactive vascular endothelium (black arrow). 7DPI: Extensive areas of pulmonary consolidation were present (black asterisk) with atypical proliferative type II pneumocyte hyperplasia (black arrow), abundant intra-alveolar macrophages degenerate neutrophils and fewer lymphocytes (black triangle). 14 DPI: There were multiple scattered areas of residual type II pneumocyte hyperplasia (black arrow), with rare clusters of macrophages, lymphocytes, and neutrophils. 28 DPI: Scattered small clusters of type II pneumocyte hyperplasia with cuboidal epithelial cells remained. Top row images at 2x magnification, bottom row 20x magnification, H&E.

#### 4 DPI

The nasal cavity still contained eosinophilic proteinaceous exudate mixed with abundant degenerate neutrophils in all infected animals and none of the uninfected animals. Olfactory epithelium was multifocally eroded with degenerate and necrotic epithelium; inflammatory cells consisted of predominantly neutrophils infiltrating into the mucosa. The tracheal submucosa only contained mild scattered infiltrates of mononuclear cells in one infected animal.

Lung changes within the medium to large airways in infected animals (Fig. 3, 4 DPI) included minimal to moderate suppurative bronchitis and bronchiolitis (8/8), bronchial and bronchiolar intraluminal neutrophilic necrotic cellular debris (8/8), bronchial epithelial syncytia (8/8), and bronchial epithelial hyperplasia (8/8). Inflammatory changes extended into the alveolar parenchyma, and included increased numbers of intra-alveolar macrophages, neutrophils, necrotic cellular debris, and eosinophilic fibrinous exudate (8/8), fibrin lining alveoli (7/8), and alveolar septal necrosis (2/8). Additionally, small foci of type II pneumocyte hyperplasia were present with large, atypical cells (8/8). Atypical cells were characterized by large, 15-20-µm diameter nuclei, with lacy chromatin and 1-3 prominent nucleoli, and moderate to large amounts of basophilic vacuolated cytoplasm. Vascular changes included reactive vascular endothelium within small to medium arteries (8/8), perivascular lymphocytic aggregates (8/8), and mild to severe vasculitis (7/8).

**Figure 4:**
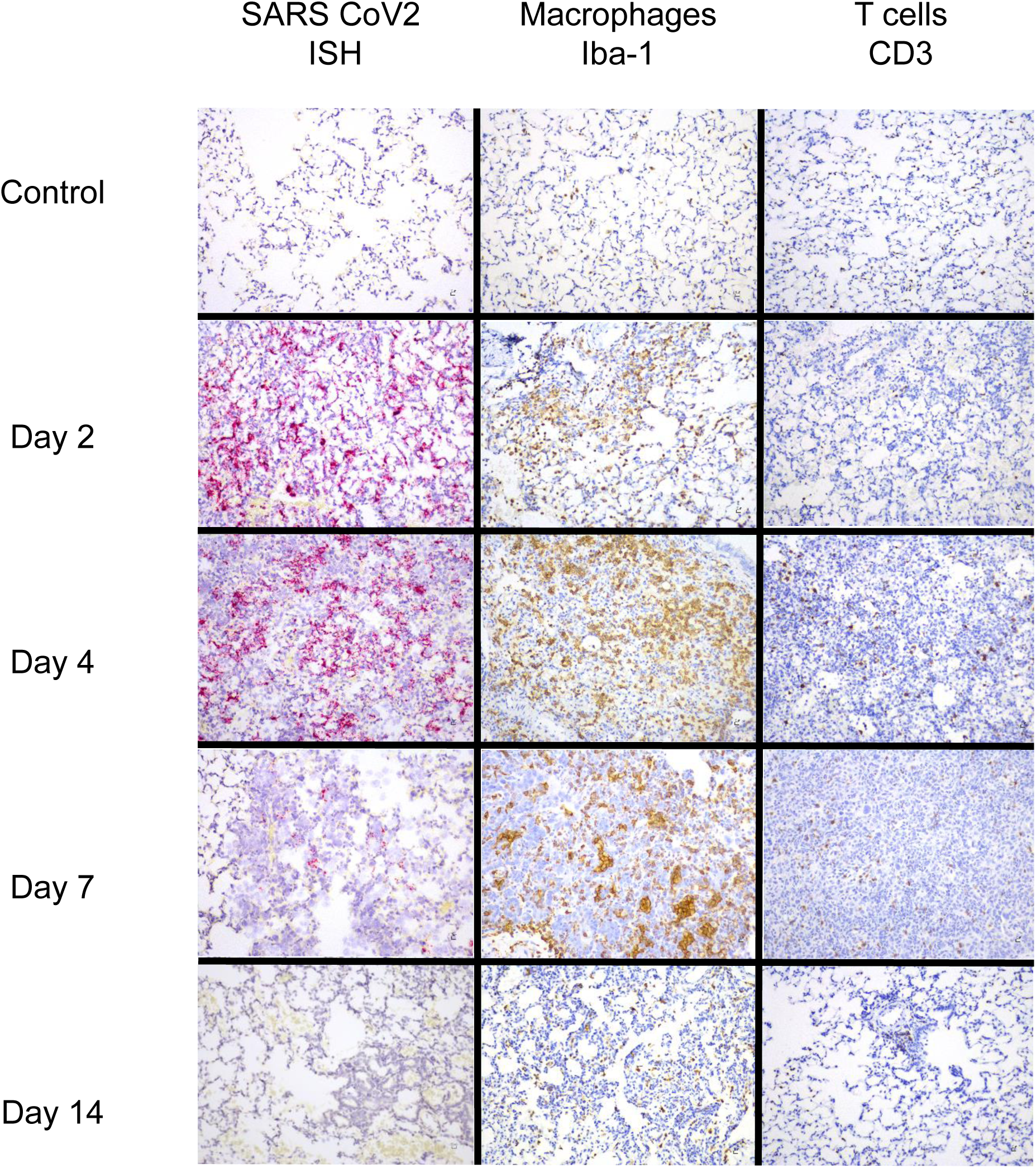
*In situ* hybridization and immunohistochemistry findings in lung. RNAscope *in situ* hybridization showed widespread, punctate staining in the lungs within alveolar epithelial cells, alveolar macrophages, bronchial and bronchiolar epithelial cells, and vascular endothelial cells at 2DPI, within alveolar epithelial cells and alveolar macrophages at 4DPI, and in one animal within alveolar epithelial cells at 7DPI. No staining was present at 14DPI (RNAscope *in situ* hybridization with red chromogen and hematoxylin counterstain). Immunostaining to detect Iba-1 revealed that macrophages comprised a major portion of the inflammatory infiltrate beginning at 2DPI and peaking at 7DPI. CD3+ lymphocytes represented a much smaller portion of the inflammatory population in the lungs but also increased from 2-7 DPI then decreased at 14 DPI. (Immunostaining with brown chromogen and hematoxylin counterstain). 20x magnification.

#### 7 DPI

The nasal cavity contained mild to moderate amounts of eosinophilic proteinaceous exudate with some degenerate neutrophils in 8/8 infected animals and 0/4 uninfected animals. At this stage, the changes in the lungs reflect a culmination of the inflammatory and reparative processes. All infected animals (8/8) developed large areas of pulmonary consolidation encompassing most of the lung (see Fig 3, 7 DPI) characterized by type II pneumocyte hyperplasia and bronchial epithelial hyperplasia that was frequently atypical, with large nuclei in variably sized cells, and frequent multinucleated cells and mitotic figures, resulting in massive thickening of the alveolar septa. Remaining alveolar air spaces in these affected areas of lung contained large numbers of macrophages with fewer degenerate neutrophils and multifocal necrotic debris. Aggregates of lymphocytes and plasma cells occasionally surrounded (8/8) and infiltrated (2/8) medium sized pulmonary arteries.

#### 14 DPI

The nasal cavity and trachea morphology were unremarkable in all infected (8/8) and control (4/4) animals. In infected animals, within the lungs (Figure 2, 14 DPI) there were scattered tufts of cuboidal, type II pneumocyte hyperplasia (8/8), rare randomly distributed clusters of macrophages that occasionally admixed with lymphocytes and neutrophils (8/8), small perivascular aggregates of lymphocytes (5/8), and multifocal small clusters of pigmented macrophages (4/8).

#### 28 DPI

The nasal cavity and trachea morphology were unremarkable in all infected (8/8) and control (4/4) animals. In infected animals, the lungs (Fig. 2, 28 DPI) contained multifocal residual areas of cuboidal, typical type II pneumocyte hyperplasia (8/8) often centered on terminal bronchioles, small clusters of pigmented macrophages (6/8), and rare aggregates of perivascular lymphocytes with pigmented macrophages (2/8). SARS-CoV-2-infected hamsters could be distinguished from uninfected animals by the presence of persistent type II pneumocyte hyperplasia in the infected animals. In contrast, lung inflammation had resolved, and fibrosis had not developed.

### Tracking SARS-CoV-2 RNA by *in situ* hybridization

RNAScope *in situ* hybridization was used to map distribution and amount of viral RNA in the lungs of infected animals throughout infection. Viral RNA expression was quantitated using QuPath pixel classification. Expression of viral RNA was significantly different over time in infected animals (one-way ANOVA, p<0.0001): 2 DPI lungs demonstrated the highest expression in alveolar epithelial cells, bronchiolar and bronchial epithelial cells, and alveolar macrophages (Figure 4). At 4 DPI, lung expression was decreased compared to 2 DPI, with positive staining for viral RNA largely restricted to the alveolar epithelial cells and alveolar macrophages in lung. At 7 DPI, lungs were largely negative; only one animal had multifocal low-level staining for viral RNA within scattered alveolar epithelial cells. In addition to lung, scattered positive staining for viral RNA was found within tracheobronchial lymph nodes and tracheal epithelium at 2 DPI (7/7 animals with tissue present), and less frequently at 4 DPI (2/5 animals with tissue present). Viral RNA was detected by qRT-PCR for up to 28 DPI in SARS-CoV-2-infected hamsters (Figure 6), suggesting that this may be the most sensitive method for viral detection, though not indicative of infectivity.

**Figure 5.**
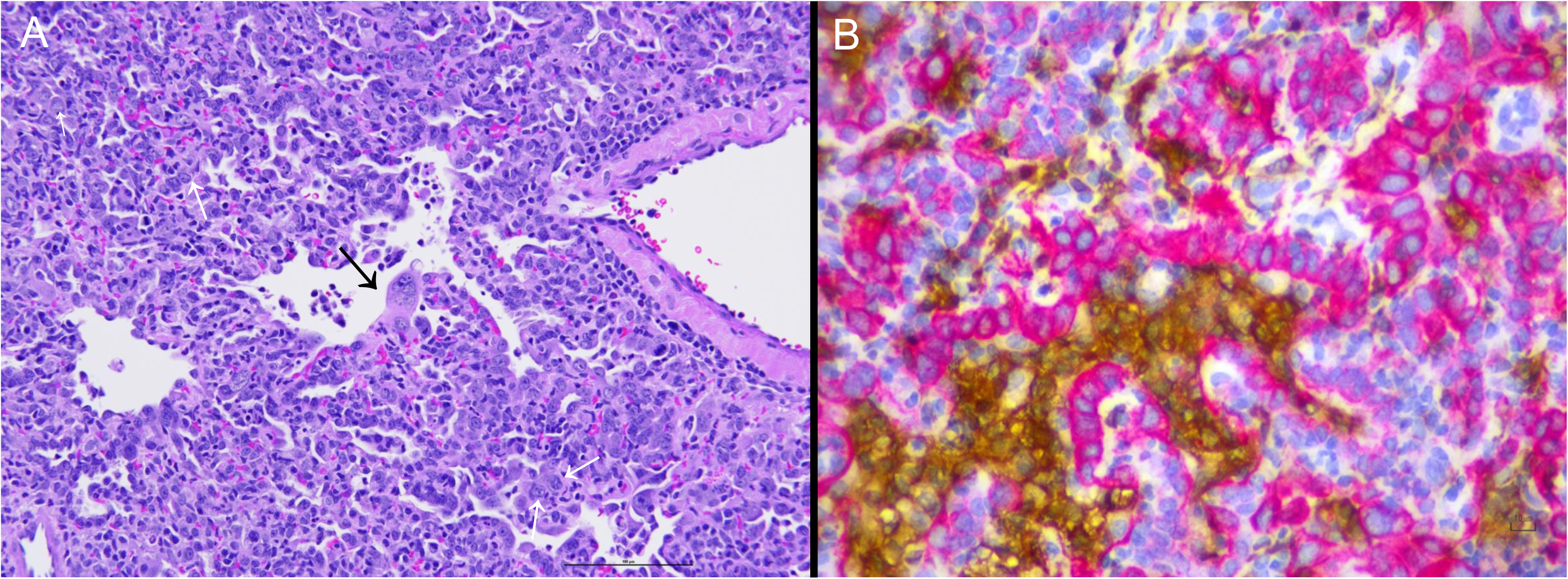
Consolidation in the lungs comprised of both large numbers of macrophages and extensive type II pneumocyte hyperplasia. A. Representative image of lung at 7DPI showing severe consolidation, including type II pneumocyte hyperplasia with large atypical epitheial cells (white arrows) and multinucleated syncytial cells (black arrow). H&E. B. Similar area of consolidation within the lungs at 7DPI with immunostaining to detect pancytokeratin (red chromogen) and Iba-1 (brown chromogen) revealed that the large, atypical cells and multinucleated cells were epithelial whereas the macrophages were within the alveolar and interstitial spaces. Hematoxylin counterstain, 40X magnification.

**Figure 6.**
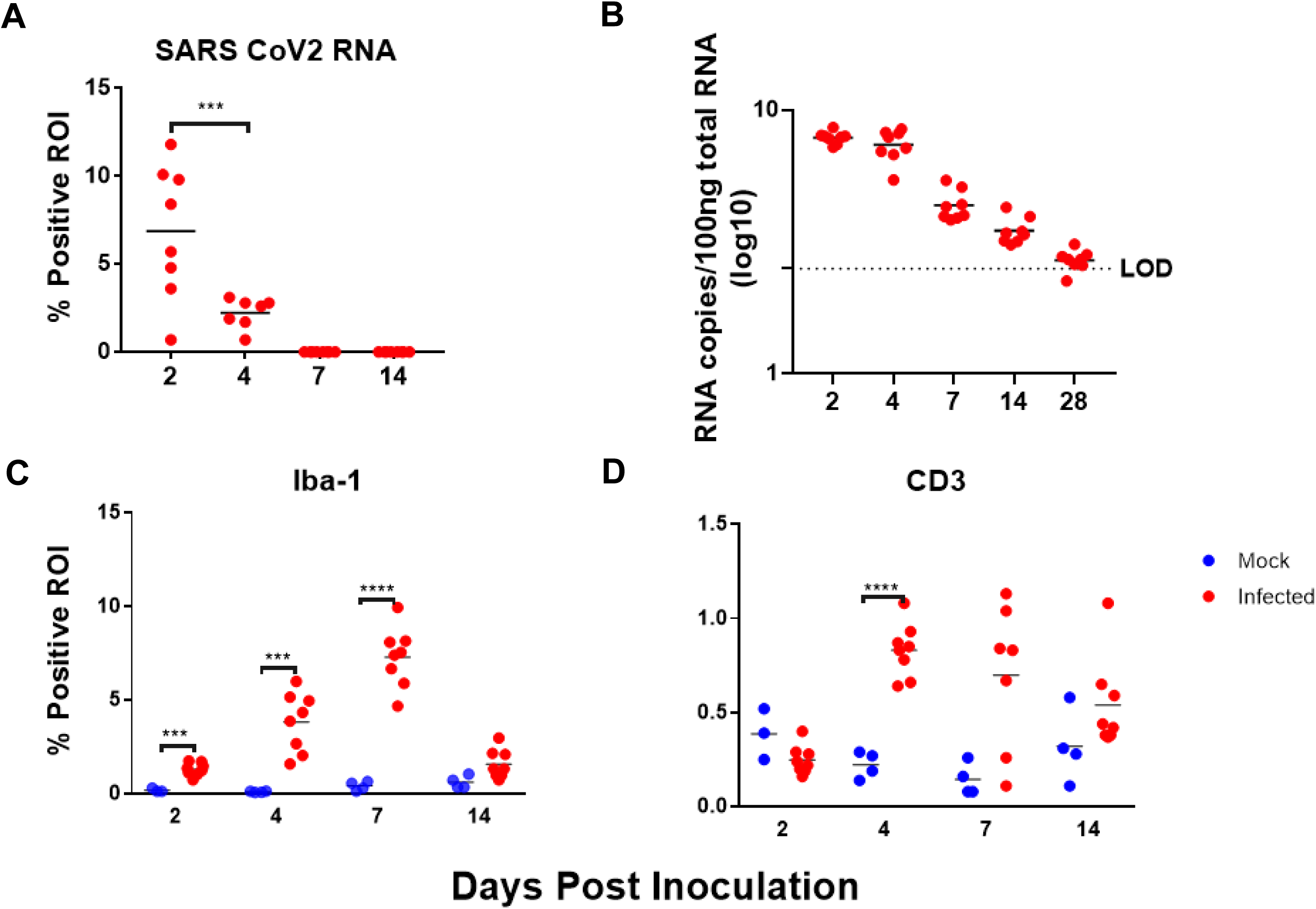
Quantitative digital image analysis of ISH and IHC staining in the lungs using QuPath analysis. A. Quantitation of staining for SARS-CoV-2 RNA within the lungs showed highest levels at 2DPI, with virus still detectable at 4 DPI. No measurable virus was detected at 7 or 14 DPI. B. Copy numbers of viral RNA within the lungs detected by qRT-PCR reached highest levels at 2 and 4 DPI, with progressively lower amounts detected at 7 DPI extending through 28 DPI. C. Quantitation of Iba-1 immunolabeling within the lungs revealed an increase in macrophages in infected animals until 7DPI, with a decrease at 14 DPI, though still above levels found in mock-inoculated animals. D. Quantitation of CD3+ immunolabeling within the lungs revealed an increase in lymphocytes until 4-7DPI. Numbers of lymphocytes were 10-fold lower than macrophages.

### Immunophenotypic Characterization of Lung Inflammation

To establish the immunophenotype and progression of inflammatory cell infiltrates in the lung, lung sections from mock-inoculated and SARS-CoV 2-infected animals were evaluated for expression of the macrophage marker Iba-1 and the T-cell marker CD3 at 2, 4, 7, and 14 DPI. Following immunostaining, QuPath was used to quantitate percent positive ROI in whole slide scanned images of the left lung lobe, the largest hamster lung lobe. Iba-1 expression was significantly different in infected animals versus uninfected animals over time (one-way ANOVA, p < 0.0001). Specifically, Iba-1 immunostaining was higher in infected animals compared with uninfected control animals at 2, 4, 7, and 14 DPI (t-tests, p < 0.001, p<0.001, P<0.0001, P= 0.04, respectively).

CD3 expression reflecting T-cell responses was more variable than Iba-1 expression. CD3 expression in infected animals was different over time (one-way ANOVA, p<0.0001). CD3 expression was significantly greater in infected animals compared with uninfected control animals at 4 and 7 DPI (t tests, p<0.0001, p=0.023, respectively). Expression of both Iba-1 and CD3 increased until 7 DPI, and then decreased at 14 DPI (Figure 4) however IBA-1 expression was much more abundant than CD3, indicating that macrophages comprised a greater portion of the inflammatory response in this model, consistent with previous reports^17^. Many of the atypical and multinucleated cells in the day 4 and day 7 animals were immunonegative for Iba-1 expression. Immunostaining for pancytokeratin revealed that numerous atypical and multinucleated cells were immunopositive, indicating that these cells were epithelial, consistent with proliferative type II pneumocytes contributing to alveolar repair (Figure 5). Double immunostaining demonstrated that both macrophages and proliferative epithelial cells were present within the densely cellular areas of the lungs of 7 DPI animals, confirming that inflammatory and reparative responses were contributing to the vast areas of consolidation at this time point.

### Quantitation of Lung Consolidation at 7 DPI

We used QuPath for digital image analysis to quantitate the percentage of lung that was affected with inflammation and reparative processes in SARS-CoV2-infected animals. The goal of this consolidation scoring approach was to encompass inflammatory cells (macrophages, neutrophils, and scattered lymphocytes), type II pneumocyte hyperplasia, necrotic cellular debris, and proteinaceous exudate while excluding artefactual collapsed or poorly insufflated regions (atelectasis). Because atelectatic areas in which alveolar septa are collapsed are common in postmortem lung examples, there was a baseline level of detection above zero in control animals. When comparing the percentage of consolidation in infected animals to the baseline level in control animals, day 7 animals had a significant increase in fold change. Infected animals at 7 DPI had a mean value of 2.978-fold change above baseline (P= 0.02, Tukey’s multiple comparisons test). The highest percentage of affected lung was 59% and occurred in a 7 DPI animal. Consolidation scores were highly correlated with IBA-1 quantitation by digital image analysis (Fig 8b; Pearson’s r = 0.84; P<0.0001).

### Flow cytometry to detect alterations in circulating cells

Fold change, relative to mock-infected hamsters at each time point, was used to assess changes in the proportions of T-lymphocyte populations circulating in blood. A significant early decrease in the proportion of CD8+ lymphocytes at 2 DPI was observed (Fig. 8A, P<0.001). No significant changes were observed in CD4+ lymphocytes (Fig. 8B, P = 0.06), however, the two 2 animals with the largest decrease in CD8+ lymphocytes also had a detectable decrease in CD4+ lymphocytes. No changes were observed in either B cells or monocyte-sized MHCII+ cells in any animals (data not shown).

**Figure 7.**
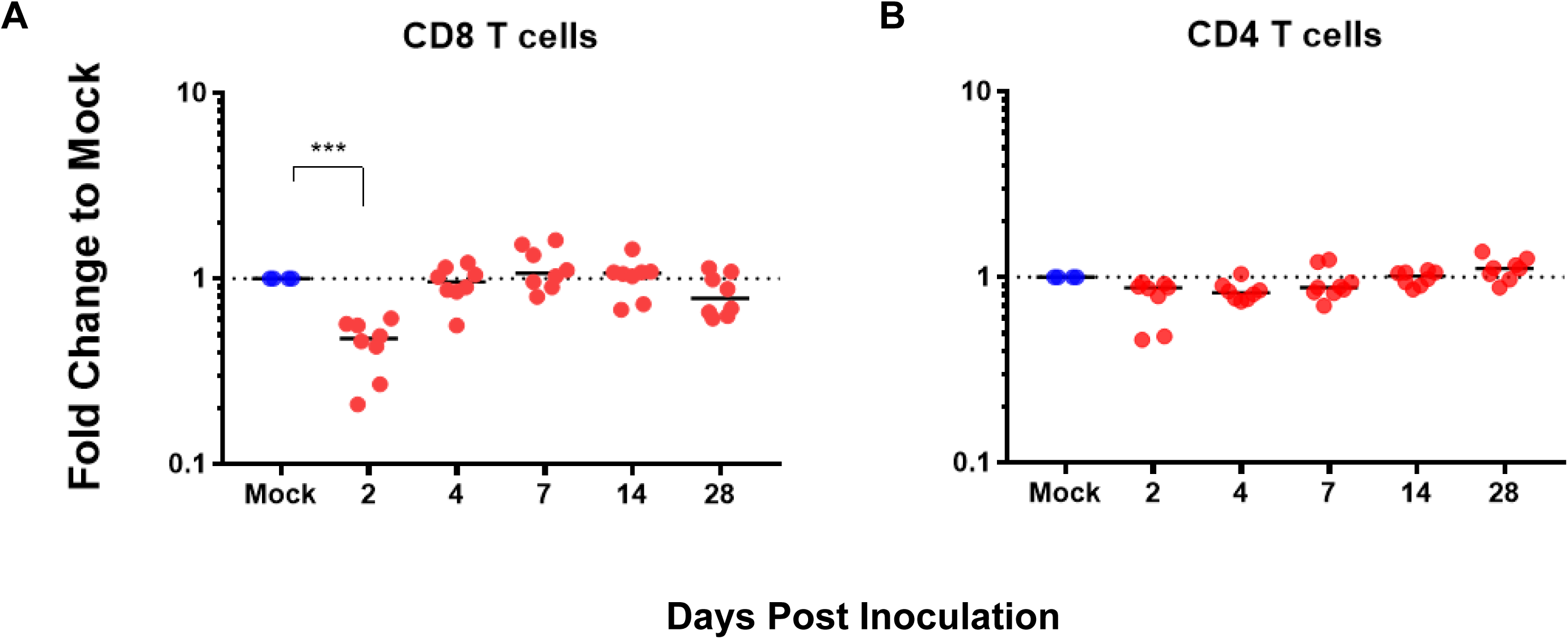
Flow cytometry of whole blood. Whole blood FACS revealed an early decrease in the proportion of circulating CD8+ lymphocytes at 2 DPI (A) while CD4 lymphocytes do not differ from baseline (B).

**Figure 8.**
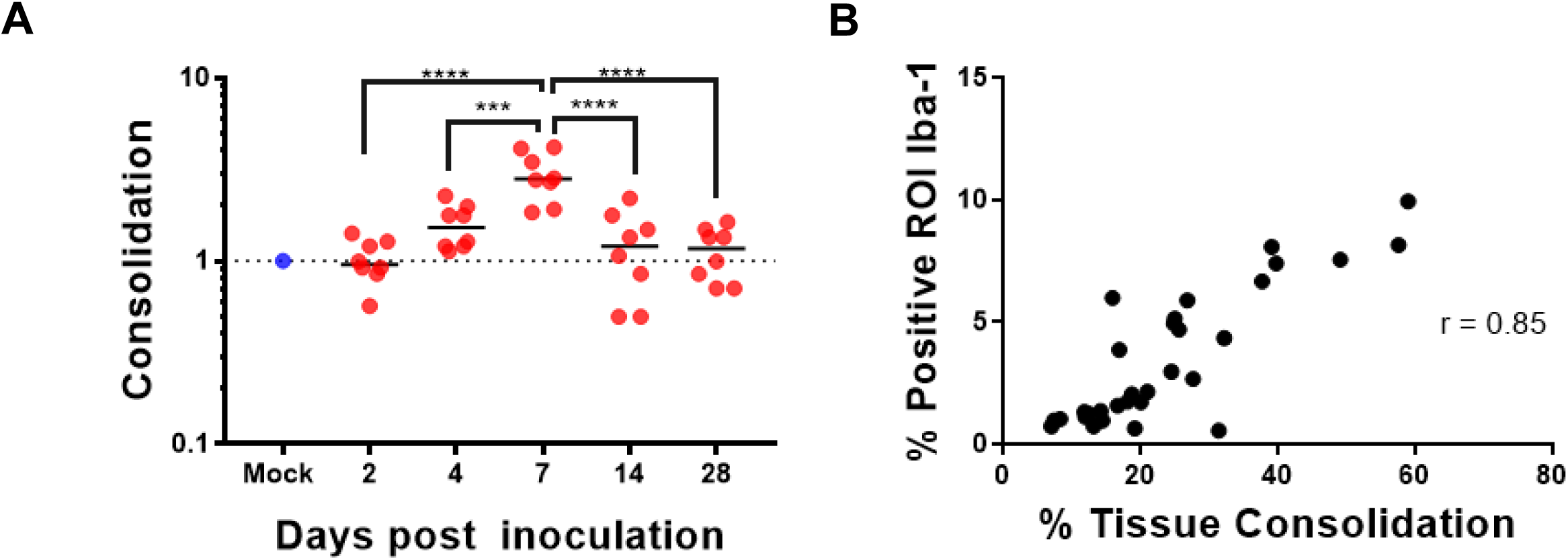

### Histopathology of Organ Systems

Minimal to mild multifocal hepatitis was observed in numerous animals. In some animals this was associated with minimal hepatocellular necrosis with few neutrophils, in others it was primarily lymphocytic. This was considered a background lesion as hepatitis was seen in control animals as well as infected animals. The bone marrow in all animals was densely cellular, with abundant myeloid cells including numerous band cells, as well as erythroid and megakaryocyte lineages. The spleen in all animals was comprised of predominantly red pulp, with no obvious signs of lymphoid depletion nor lymphoid hyperplasia. Abdominal lymph nodes occasionally contained germinal centers in control and infected animals. Incidental findings included rare protozoa associated with superficial gastric mucosa (12/48), morphologically suggestive of *Cryptosporidium* spp, with infrequent and mild hyperplasia of the affected gastric mucosa. No significant changes were noted within the remaining gastrointestinal tract, including the oral cavity, esophagus, small intestine, cecum, and colon. Kidneys in animals across all groups (29/45) had minimal protein and mineral within the tubules. Adrenal glands within hamsters of all groups had variable numbers of pigmented cells within the cortex, which has been reported to be a change in aged hamsters^30^. Other organs that were examined without significant changes on histopathology included: brain, heart, gallbladder, male and female reproductive organs, urinary bladder, salivary glands, bone, haired skin, skeletal muscle, and decalcified cross sections of the head.

## Discussion

In this study, we conducted a longitudinal, in-depth comprehensive histopathological analysis of all tissues and organ systems within male and female golden Syrian hamsters inoculated intranasally with SARS-CoV-2, spanning 2 to 28 days post-inoculation. Additionally, we characterized the immunophenotype of the inflammatory infiltrates within the lungs across all stages of disease and assessed viral load within the lungs via *in situ* hybridization, virus titration, and qRT-PCR for viral RNA. We quantified these morphologic findings using QuPath, an open-source digital image analysis platform applied to analyze an entire lung lobe imaged via whole slide scanning. This objective approach provided robust outcome measures for IHC, ISH, and consolidation, enhancing rigor of this COVID-19 model.

The main lesion type within the lungs was proliferative bronchointerstitial pneumonia, which started as acute necrosis of alveolar septa and airway epithelium coincident with SARS-CoV-2 replication. Cellular immune responses were first dominated by recruited neutrophils but switched to a macrophage-dominant phenotype accompanied by a robust atypical type II pneumocyte reparative response beginning by day 4 and peaking at day 7 post-inoculation, resembling other reports of disease timing in this animal model ^13–18^. Pulmonary pathology findings closely emulated changes reported in humans which include diffuse alveolar damage, intra-alveolar macrophage infiltration, and type II pneumocyte hyperplasia^31^. Atypical proliferation of type II pneumocytes in the hamster model during the reparative phase of disease is inconsistently reported but was a very consistent and prominent finding in this study^14^. The striking proliferation was more robust than what has been reported in humans with frequent anisocytosis and anisokaryosis and numerous mitotic figures. Type II pneumocyte hyperplasia in hamsters has been reported in other experimentally induced respiratory viral infections^32,33^, but is not usually this florid. Hamsters were evaluated from initial stages of infection at progressive time points, whereas reports of human COVID-19 lung pathology are cross-sectional, typically at autopsy representing the most severe disease, the timing of lung sampling may explain this disparity.

Vascular changes are consistently described in human SARS-CoV-2 infection and are thought to be a key factor in the pathogenesis of severely affected individuals^4,31^. Within the lungs, vascular changes in humans include endothelial damage, thrombosis, microangiopathy, congestion, and angiogenesis^4^. In this study, within the pulmonary vasculature, there was prominent endothelial hypertrophy in days 2-7, as well as overt vasculitis in small and medium-sized arteries predominantly at day 4. Further exploration and characterization of these changes in future studies is warranted given the importance that these factors may play in human disease. Interstitial fibrosis is a chronic sequela to SARS-CoV-2 infection in some individuals^3,31^. Interestingly, while interstitial fibrosis was not observed in this study, there was residual type II pneumocyte hyperplasia at day 14 and 28 frequently present adjacent to terminal bronchioles, which has not been reported in COVID-19 animal models or in human COVID-19 cases likely because these changes develop during the reparative stage time points when post-mortem samples are not typically available.

In this model, a significant decrease in the proportion of CD8+ lymphocytes in peripheral blood of infected hamsters was detected at day 2 DPI. Further investigation is warranted to characterize this change to determine if this is representative of a CD8- dominant lymphopenia, as this could have important implications for pathogenesis and may be representative of human disease ^24,34–37^. In human COVID-19 patients, CD8- dominant lymphopenia has been consistently documented and critical patients have more severe lymphopenia than individuals with mild cases^35,37,38^. The mechanisms underlying this change are unclear. In this study we did not observe destructive changes in the thymus, spleen, lymph nodes or gut-associated lymphoid tissue suggestive of lymphoid depletion.

Overall, hamsters developed consistent pulmonary disease associated with SARS-CoV- 2 infection progressing though distinct stages: acute viral replication and cell necrosis accompanied by infiltrating neutrophils, a transition to macrophage-dominant inflammation with control of viral replication and robust reparative epithelial responses when lung consolidation were most severe 7 DPI, and resolution of inflammation with residual evidence of epithelial repair still evident 28 DPI with SARS-CoV-2. Hamsters are susceptible to infection with an intact immune system, in contrast to mouse model systems in which the mouse or virus needs to be altered in order to study this pathogen and disease. Hamsters also have the advantage of being a small animal model, which are less expensive and easier to maintain than larger animals such as nonhuman primates. The consistency of disease in this model further emphasizes the need to develop additional antibodies and reagents to characterize immunopathologic changes.

This study provides detailed and objectively quantified longitudinal histopathological analysis of the respiratory system, and describes associated changes observed in all tissues and organ systems. These comprehensive and integrated findings on disease progression and resolution serve as a baseline of SAS-CoV-2 infection outcome measures in this model and will be valuable for determining the efficacy of therapeutic and preventive interventions for COVID-19.

## Acknowledgements

Additional Johns Hopkins COVID-19 Hamster Study Group members include: Michael J. Betenbaugh, Bess Carlson, Natalie Castell, Jennie Ruelas Castillo, Kelly Flavahan, Eric K. Hutchinson, Kirsten Littlefield, Monika M. Looney, Maggie Lowman, Natalia Majewski, Amanda Maxwell, Filipa Mota, Alice L. Mueller, Alvaro A. Ordonez, Lisa Pieterse, Darla Quijada, Camilo A. Ruiz-Bedoya, Mitchel Stover, Rachel Vistein, Melissa Wood and Cynthia A. Zahnow.

AP would like to dedicate this manuscript to the memory of R. Mark Buller, whose collaborations on the golden Syrian hamster model for SARS-CoV infection formed the basis for this study.

## Supplemental QuPath Script Information

Note: Annotation of lung sections was performed individually using the “wand” tool.

### IBA-1 detection and quantification

setImageType(’BRIGHTFIELD_H_DAB’);

setColorDeconvolutionStains(’{“Name” : “H-DAB default”, “Stain 1” : “Hematoxylin”, “Values 1” : “0.65111 0.70119 0.29049 “, “Stain 2” : “DAB”, “Values 2” : “0.26917 0.56824 0.77759 “, “Background” : “ 255 255 255 “”’);

addPixelClassifierMeasurements(“IBA1_Lung_1”, “IBA1_Lung_1”)

### CD3 detection and quantification

setImageType(’BRIGHTFIELD_H_DAB’);

setColorDeconvolutionStains(’{“Name” : “H-DAB default”, “Stain 1” : “Hematoxylin”, “Values 1” : “0.65111 0.70119 0.29049 “, “Stain 2” : “DAB”, “Values 2” : “0.26917 0.56824 0.77759 “, “Background” : “ 255 255 255 “”’);

selectAnnotations();

addPixelClassifierMeasurements(“CD3_5”, “CD3_5”)

### ISH detection and quantification

setImageType(’BRIGHTFIELD_H_DAB’);

setColorDeconvolutionStains(’{“Name” : “H-DAB default”, “Stain 1” : “Hematoxylin”, “Values 1” : “0.65111 0.70119 0.29049 “, “Stain 2” : “DAB”, “Values 2” : “0.26917 0.56824 0.77759 “, “Background” : “ 255 255 255 “”’);

selectAnnotations(); addPixelClassifierMeasurements(“SARS_4”, “SARS_4”)

### Consolidation

setImageType(’BRIGHTFIELD_H_E’);

setColorDeconvolutionStains(’{“Name” : “H&E default”, “Stain 1” : “Hematoxylin”, “Values 1” : “0.65111 0.70119 0.29049 “, “Stain 2” : “Eosin”, “Values 2” : “0.2159 0.8012 0.5581 “, “Background” : “ 255 255 255 “”’);

selectAnnotations();

runPlugin(’qupath.imagej.superpixels.DoGSuperpixelsPlugin’, ’{“downsampleFactor”: 8.0, “sigmaMicrons”: 5.0, “minThreshold”: 15.0, “maxThreshold”: 200.0, “noiseThreshold”: 0.5}’);

selectDetections();

runPlugin(’qupath.lib.algorithms.IntensityFeaturesPlugin’, ’{“pixelSizeMicrons”: 1.0, “region”: “ROI”, “tileSizeMicrons”: 25.0, “colorOD”: false, “colorStain1”: true, “colorStain2”: true, “colorStain3”: true, “colorRed”: false, “colorGreen”: false, “colorBlue”: false, “colorHue”: false, “colorSaturation”: false, “colorBrightness”: false, “doMean”: true, “doStdDev”: false, “doMinMax”: true, “doMedian”: false, “doHaralick”: true, “haralickDistance”: 1, “haralickBins”: 32}’);

runObjectClassifier(“D7_8”);

**Supplemental figure 1.**
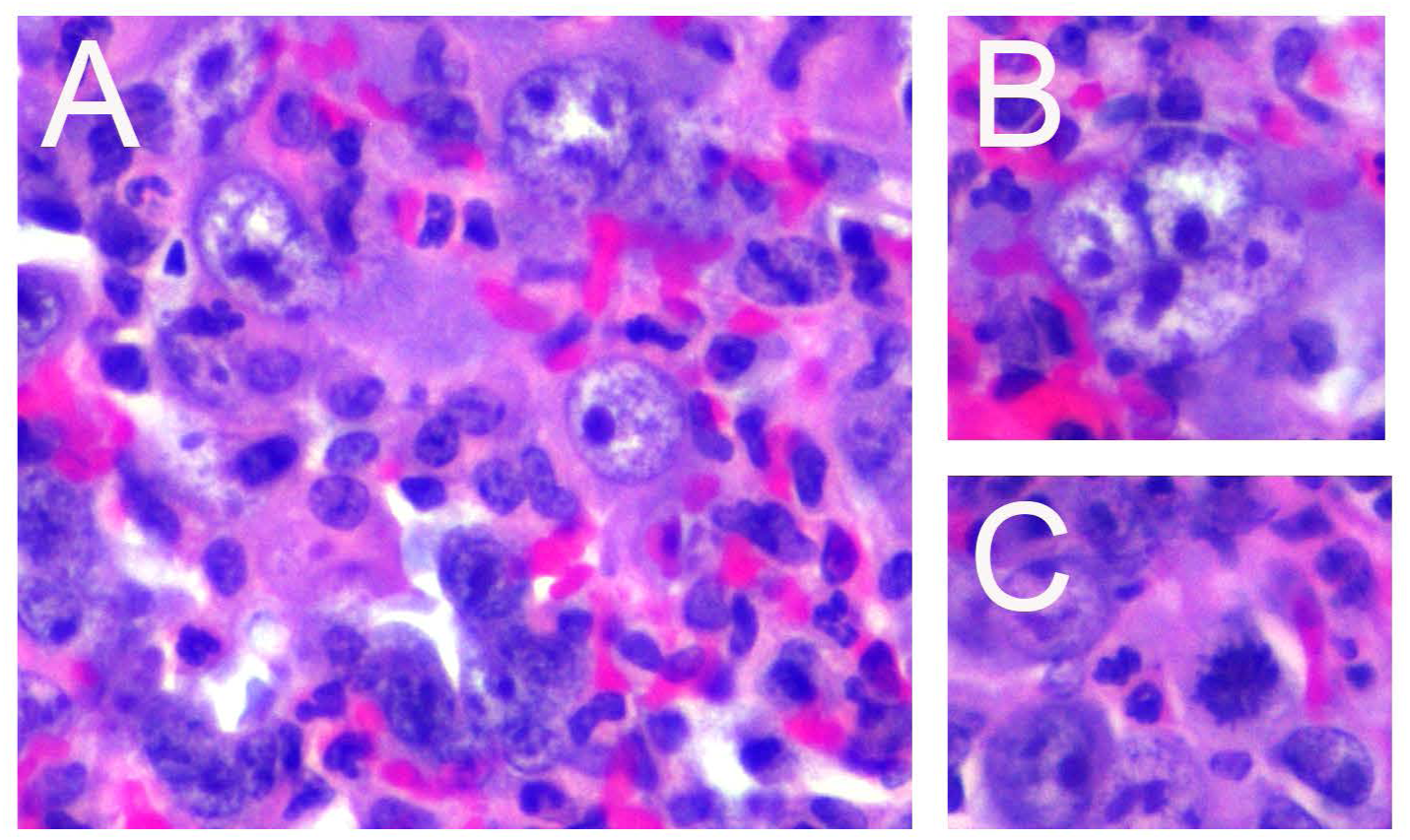
Additional images of lung pathology at 7 DPI. A. Many large, atypical cells were present lining the alveolar septa that contained large nuclei with prominent nucleoli and lacy chromatin. B. Frequent multinucleated epithelial cells were present throughout affected areas of lung. C. Numerous mitotic figures were present within areas of type II pneumocyte hyperplasia indicating a robust reparative response. 40X magnification, H&E.

**Supplemental figure 2.**
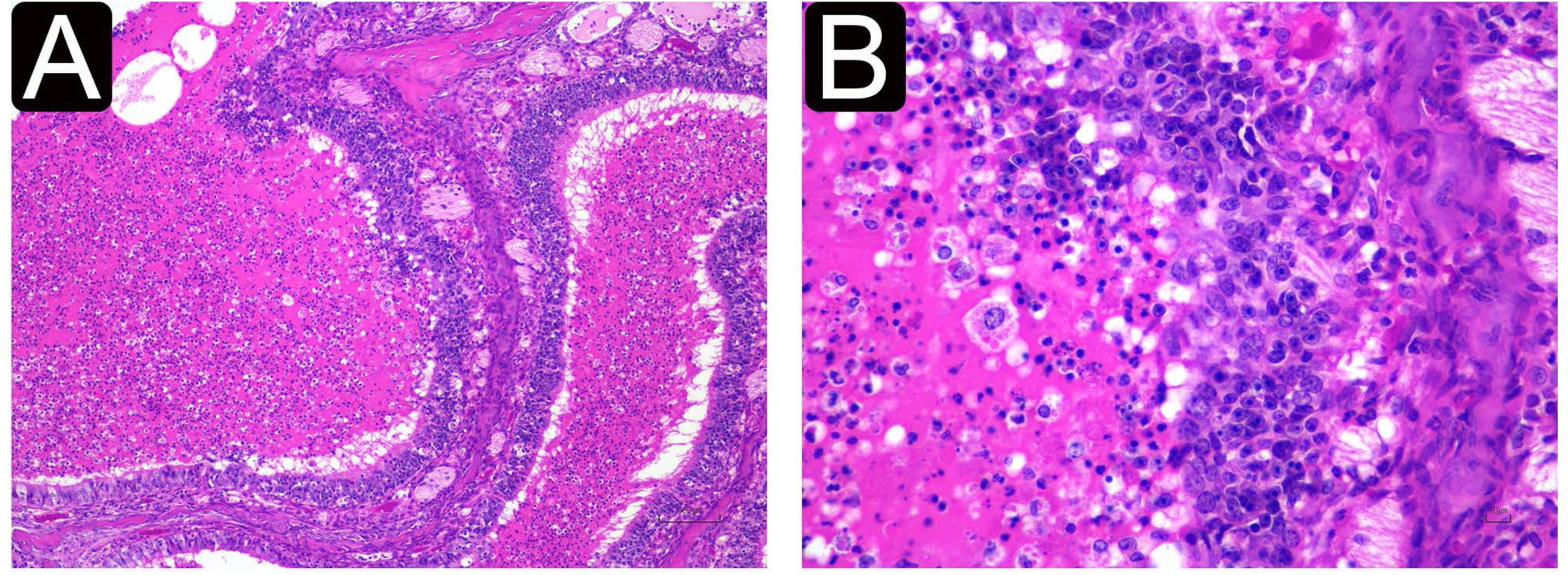
Pathology of the nasal cavity in SARS-CoV-2-infected hamsters. At 2 and 4 DPI, the nasal cavity of infected animals contained abundant suppurative exudate admixed with proteinaceous material. Inflammatory cells infiltrated into the nasal mucosal epithelium, and there were multiple areas of epithelial erosion and ulceration. A. 2x magnification. B. 20x magnification.

**Supplemental figure 3.**
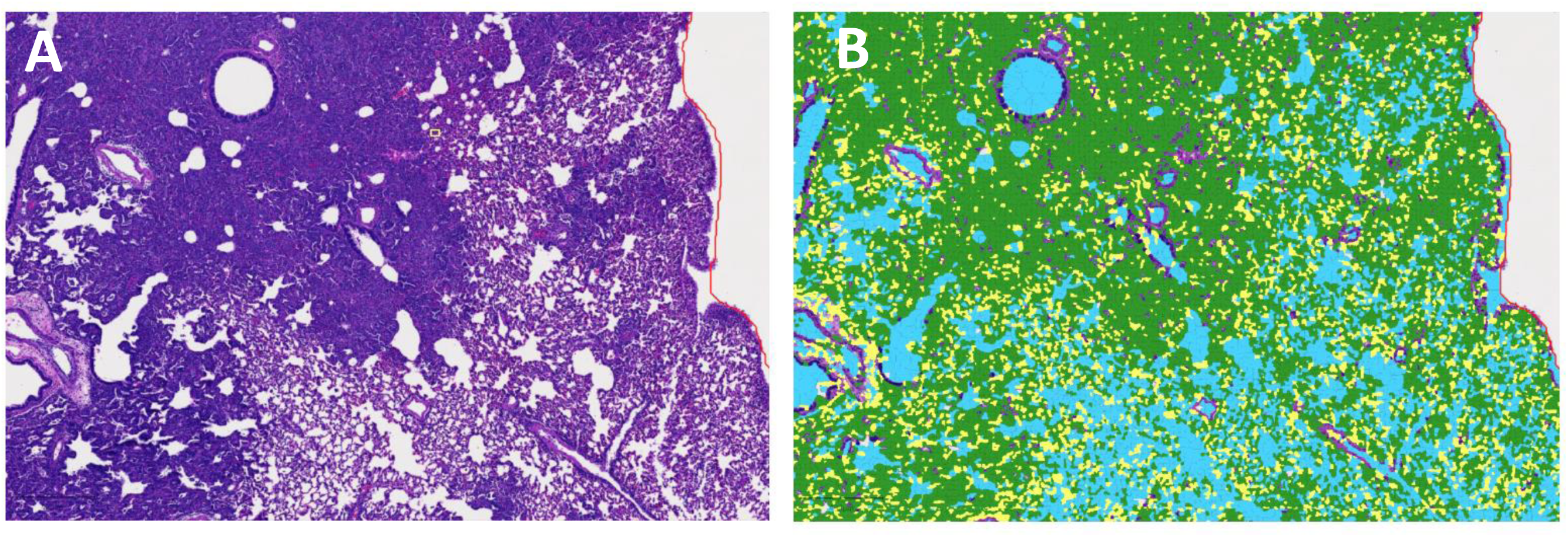
Quantitation of consolidation of the lungs at 7 DPI using QuPath. A. Representative image of SARS-CoV-2 infected animal at 7 DPI showing multiple coalescing areas of consolidation. B. Quantitation of affected area using QuPath- generated superpixel classified as either “consolidation” (green), “unaffected” (blue), or “atelectasis” (yellow). Areas where tissue is visible (bronchiolar epithelium, blood vessels), represent superpixels classfied as “ignore”. C. Quantitation of consolidated area. Infected animals had significantly higher percentage of tissue consolidation compared to mock animals (P = 0.0007, unpaired t-test)

**Supplemental figure 4.**
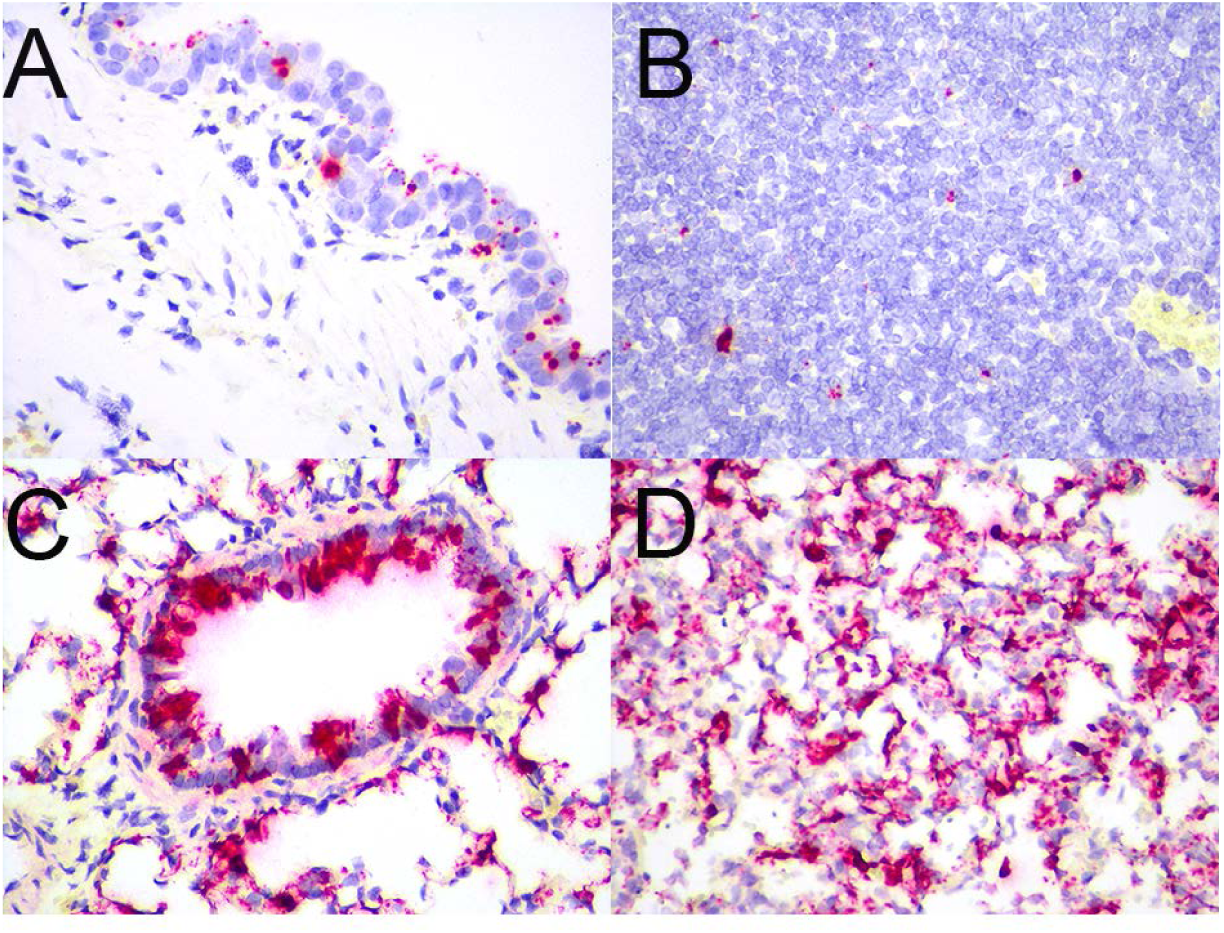
*In situ* hybridization of SARS Co-V 2 at 2DPI. SARS CO-V 2 RNA (red chromogen) is detectable within the tracheal epithelium (A), bronchial lymph node (B), bronchiolar epithelium (C), and within alveolar epithelial cells and macrophages (D). 40X magnification, hematoxylin counterstain.

## References

1. Ren L-L, Wang Y-M, Wu Z-Q, Xiang Z-C, Guo L, Xu T, Jiang Y-Z, Xiong Y, Li Y-J, Li X-W, Li H, Fan G-H, Gu X-Y, Xiao Y, Gao H, Xu J-Y, Yang F, Wang X-M, Wu C, Chen L, Liu Y-W, Liu B, Yang J, Wang X-R, Dong J, Li L, Huang C-L, Zhao J-P, Hu Y, Cheng Z-S, Liu L-L, Qian Z-H, Qin C, Jin Q, Cao B, Wang J-W: Identification of a novel coronavirus causing severe pneumonia in human: a descriptive study. Chin Med J 2020, 133:1015–1024.

2. World Health Organization: Coronavirus Disease (Covid-19) Pandemic [Internet]. 2020 [cited 2021 Feb 17],. Available from: https://www.who.int/emergencies/diseases/novel-coronavirus-2019

3. Bösmüller H, Matter M, Fend F, Tzankov A: The pulmonary pathology of COVID-19. Virchows Archiv 2021,.

4. Ackermann M, Verleden SE, Kuehnel M, Haverich A, Welte T, Laenger F, Vanstapel A, Werlein C, Stark H, Tzankov A, Li WW, Li VW, Mentzer SJ, Jonigk D: Pulmonary Vascular Endothelialitis, Thrombosis, and Angiogenesis in Covid-19. N Engl J Med 2020, 383:120–128.

5. Menter T, Haslbauer JD, Nienhold R, Savic S, Hopfer H, Deigendesch N, Frank S, Turek D, Willi N, Pargger H, Bassetti S, Leuppi JD, Cathomas G, Tolnay M, Mertz KD, Tzankov A: Postmortem examination of COVID-19 patients reveals diffuse alveolar damage with severe capillary congestion and variegated findings in lungs and other organs suggesting vascular dysfunction. Histopathology 2020, 77:198– 209.

6. Bösmüller H, Traxler S, Bitzer M, Häberle H, Raiser W, Nann D, Frauenfeld L, Vogelsberg A, Klingel K, Fend F: The evolution of pulmonary pathology in fatal COVID-19 disease: an autopsy study with clinical correlation. Virchows Arch 2020, 477:349–357.

7. Lax SF, Skok K, Zechner P, Kessler HH, Kaufmann N, Koelblinger C, Vander K, Bargfrieder U, Trauner M: Pulmonary Arterial Thrombosis in COVID-19 With Fatal Outcome : Results From a Prospective, Single-Center, Clinicopathologic Case Series. Ann Intern Med 2020, 173:350–361.

8. Carsana L, Sonzogni A, Nasr A, Rossi RS, Pellegrinelli A, Zerbi P, Rech R, Colombo R, Antinori S, Corbellino M, Galli M, Catena E, Tosoni A, Gianatti A, Nebuloni M: Pulmonary post-mortem findings in a series of COVID-19 cases from northern Italy: a two-centre descriptive study. Lancet Infect Dis 2020, 20:1135–1140.

9. Winkler ES, Bailey AL, Kafai NM, Nair S, McCune BT, Yu J, Fox JM, Chen RE, Earnest JT, Keeler SP, Ritter JH, Kang L-I, Dort S, Robichaud A, Head R, Holtzman MJ, Diamond MS: SARS-CoV-2 infection of human ACE2-transgenic mice causes severe lung inflammation and impaired function. Nat Immunol 2020, 21:1327–1335.

10. Winkler ES, Bailey AL, Kafai NM, Nair S, McCune BT, Yu J, Fox JM, Chen RE, Earnest JT, Keeler SP, Ritter JH, Kang L-I, Dort S, Robichaud A, Head R, Holtzman MJ, Diamond MS: SARS-CoV-2 infection in the lungs of human ACE2 transgenic mice causes severe inflammation, immune cell infiltration, and compromised respiratory function. BioRxiv 2020,.

11. Sun S-H, Chen Q, Gu H-J, Yang G, Wang Y-X, Huang X-Y, Liu S-S, Zhang N-N, Li X-F, Xiong R, Guo Y, Deng Y-Q, Huang W-J, Liu Q, Liu Q-M, Shen Y-L, Zhou Y, Yang X, Zhao T-Y, Fan C-F, Zhou Y-S, Qin C-F, Wang Y-C: A Mouse Model of SARS-CoV-2 Infection and Pathogenesis. Cell Host Microbe 2020, 28:124–133.e4.

12. Abdel-Moneim AS, Abdelwhab EM: Evidence for SARS-CoV-2 Infection of Animal Hosts. Pathogens 2020, 9.

13. Sia SF, Yan L-M, Chin AWH, Fung K, Choy K-T, Wong AYL, Kaewpreedee P, Perera RAPM, Poon LLM, Nicholls JM, Peiris M, Yen H-L: Pathogenesis and transmission of SARS-CoV-2 in golden hamsters. Nature 2020, 583:834–838.

14. Chan JF-W, Zhang AJ, Yuan S, Poon VK-M, Chan CC-S, Lee AC-Y, Chan W-M, Fan Z, Tsoi H-W, Wen L, Liang R, Cao J, Chen Y, Tang K, Luo C, Cai J-P, Kok K-H, Chu H, Chan K-H, Sridhar S, Chen Z, Chen H, To KK-W, Yuen K-Y: Simulation of the Clinical and Pathological Manifestations of Coronavirus Disease 2019 (COVID-19) in a Golden Syrian Hamster Model: Implications for Disease Pathogenesis and Transmissibility. Clin Infect Dis 2020, 71:2428–2446.

15. Imai M, Iwatsuki-Horimoto K, Hatta M, Loeber S, Halfmann PJ, Nakajima N, Watanabe T, Ujie M, Takahashi K, Ito M, Yamada S, Fan S, Chiba S, Kuroda M, Guan L, Takada K, Armbrust T, Balogh A, Furusawa Y, Okuda M, Ueki H, Yasuhara A, Sakai-Tagawa Y, Lopes TJS, Kiso M, Yamayoshi S, Kinoshita N, Ohmagari N, Hattori S-I, Takeda M, Mitsuya H, Krammer F, Suzuki T, Kawaoka Y: Syrian hamsters as a small animal model for SARS-CoV-2 infection and countermeasure development. Proc Natl Acad Sci USA 2020, 117:16587–16595.

16. Osterrieder N, Bertzbach LD, Dietert K, Abdelgawad A, Vladimirova D, Kunec D, Hoffmann D, Beer M, Gruber AD, Trimpert J: Age-dependent progression of SARS-CoV-2 infection in Syrian hamsters. BioRxiv 2020,.

17. Tostanoski LH, Wegmann F, Martinot AJ, Loos C, McMahan K, Mercado NB, Yu J, Chan CN, Bondoc S, Starke CE, Nekorchuk M, Busman-Sahay K, Piedra-Mora C, Wrijil LM, Ducat S, Custers J, Atyeo C, Fischinger S, Burke JS, Feldman J, Hauser BM, Caradonna TM, Bondzie EA, Dagotto G, Gebre MS, Jacob-Dolan C, Lin Z, Mahrokhian SH, Nampanya F, Nityanandam R, Pessaint L, Porto M, Ali V, Benetiene D, Tevi K, Andersen H, Lewis MG, Schmidt AG, Lauffenburger DA, Alter G, Estes JD, Schuitemaker H, Zahn R, Barouch DH: Ad26 vaccine protects against SARS-CoV-2 severe clinical disease in hamsters. Nat Med 2020, 26:1694–1700.

18. Kreye J, Reincke SM, Kornau H-C, Sánchez-Sendin E, Corman VM, Liu H, Yuan M, Wu NC, Zhu X, Lee C-CD, Trimpert J, Höltje M, Dietert K, Stöffler L, von Wardenburg N, van Hoof S, Homeyer MA, Hoffmann J, Abdelgawad A, Gruber AD, Bertzbach LD, Vladimirova D, Li LY, Barthel PC, Skriner K, Hocke AC, Hippenstiel S, Witzenrath M, Suttorp N, Kurth F, Franke C, Endres M, Schmitz D, Jeworowski LM, Richter A, Schmidt ML, Schwarz T, Müller MA, Drosten C, Wendisch D, Sander LE, Osterrieder N, Wilson IA, Prüss H: A Therapeutic Non-self-reactive SARS-CoV-2 Antibody Protects from Lung Pathology in a COVID-19 Hamster Model. Cell 2020, 183:1058–1069.e19.

19. Munster VJ, Feldmann F, Williamson BN, van Doremalen N, Pérez-Pérez L, Schulz J, Meade-White K, Okumura A, Callison J, Brumbaugh B, Avanzato VA, Rosenke R, Hanley PW, Saturday G, Scott D, Fischer ER, de Wit E: Respiratory disease in rhesus macaques inoculated with SARS-CoV-2. Nature 2020, 585:268–272.

20. Baum A, Ajithdoss D, Copin R, Zhou A, Lanza K, Negron N, Ni M, Wei Y, Mohammadi K, Musser B, Atwal GS, Oyejide A, Goez-Gazi Y, Dutton J, Clemmons E, Staples HM, Bartley C, Klaffke B, Alfson K, Gazi M, Gonzalez O, Dick E, Carrion R, Pessaint L, Porto M, Cook A, Brown R, Ali V, Greenhouse J, Taylor T, Andersen H, Lewis MG, Stahl N, Murphy AJ, Yancopoulos GD, Kyratsous CA: REGN-COV2 antibodies prevent and treat SARS-CoV-2 infection in rhesus macaques and hamsters. Science 2020, 370:1110–1115.

21. Kim Y-I, Kim S-G, Kim S-M, Kim E-H, Park S-J, Yu K-M, Chang J-H, Kim EJ, Lee S, Casel MAB, Um J, Song M-S, Jeong HW, Lai VD, Kim Y, Chin BS, Park J-S, Chung K-H, Foo S-S, Poo H, Mo I-P, Lee O-J, Webby RJ, Jung JU, Choi YK: Infection and Rapid Transmission of SARS-CoV-2 in Ferrets. Cell Host Microbe 2020, 27:704–709.e2.

22. Muñoz-Fontela C, Dowling WE, Funnell SGP, Gsell P-S, Riveros-Balta AX, Albrecht RA, Andersen H, Baric RS, Carroll MW, Cavaleri M, Qin C, Crozier I, Dallmeier K, de Waal L, de Wit E, Delang L, Dohm E, Duprex WP, Falzarano D, Finch CL, Frieman MB, Graham BS, Gralinski LE, Guilfoyle K, Haagmans BL, Hamilton GA, Hartman AL, Herfst S, Kaptein SJF, Klimstra WB, Knezevic I, Krause PR, Kuhn JH, Le Grand R, Lewis MG, Liu W-C, Maisonnasse P, McElroy AK, Munster V, Oreshkova N, Rasmussen AL, Rocha-Pereira J, Rockx B, Rodríguez E, Rogers TF, Salguero FJ, Schotsaert M, Stittelaar KJ, Thibaut HJ, Tseng C-T, Vergara-Alert J, Beer M, Brasel T, Chan JFW, García-Sastre A, Neyts J, Perlman S, Reed DS, Richt JA, Roy CJ, Segalés J, Vasan SS, Henao-Restrepo AM, Barouch DH: Animal models for COVID-19. Nature 2020, 586:509–515.

23. Zeiss CJ, Compton S, Veenhuis RT: Animal Models of COVID-19. I. Comparative Virology and Disease Pathogenesis. ILAR J 2021,.

24. Veenhuis RT, Zeiss CJ: Animal Models of COVID-19 II. Comparative Immunology. ILAR J 2021,.

25. Rogers TF, Zhao F, Huang D, Beutler N, Burns A, He W-T, Limbo O, Smith C, Song G, Woehl J, Yang L, Abbott RK, Callaghan S, Garcia E, Hurtado J, Parren M, Peng L, Ramirez S, Ricketts J, Ricciardi MJ, Rawlings SA, Wu NC, Yuan M, Smith DM, Nemazee D, Teijaro JR, Voss JE, Wilson IA, Andrabi R, Briney B, Landais E, Sok D, Jardine JG, Burton DR: Isolation of potent SARS-CoV-2 neutralizing antibodies and protection from disease in a small animal model. Science 2020, 369:956–963.

26. Bankhead P, Loughrey MB, Fernández JA, Dombrowski Y, McArt DG, Dunne PD, McQuaid S, Gray RT, Murray LJ, Coleman HG, James JA, Salto-Tellez M, Hamilton PW: QuPath: Open source software for digital pathology image analysis. Sci Rep 2017, 7:16878.

27. Morriss NJ, Conley GM, Ospina SM, Meehan Iii WP, Qiu J, Mannix R: Automated quantification of immunohistochemical staining of large animal brain tissue using qupath software. Neuroscience 2020, 429:235–244.

28. Dhakal S, Ruiz-Bedoya CA, Zhou R, Creisher P, Villano J, Littlefield K, Castillo J, Marinho P, Jedlicka A, Ordonez A, Majewska N, Betenbaugh M, Flavahan K, Mueller A, Looney M, Quijada D, Mota F, Beck SE, Brockhurst JK, Braxton A, Castell N, D’Alessio F, Metcalf Pate KA, Karakousis PC, Mankowski JL, Pekosz A, Jain SK, Klein SL: Sex differences in lung imaging and SARS-CoV-2 antibody responses in a COVID-19 golden Syrian hamster model. BioRxiv 2021, .

29. Gniazdowski V, Morris CP, Wohl S, Mehoke T, Ramakrishnan S, Thielen P, Powell H, Smith B, Armstrong DT, Herrera M, Reifsnyder C, Sevdali M, Carroll KC, Pekosz A, Mostafa HH: Repeat COVID-19 Molecular Testing: Correlation of SARS-CoV-2 Culture with Molecular Assays and Cycle Thresholds. Clin Infect Dis 2020,.

30. Meyers MW, Charipper HA: A histological and cytological study of the adrenal gland of the golden hamster (Cricetus auratus) in relation to age. Anat Rec 1956, 124:1–25.

31. Polak SB, Van Gool IC, Cohen D, von der Thüsen JH, van Paassen J: A systematic review of pathological findings in COVID-19: a pathophysiological timeline and possible mechanisms of disease progression. Mod Pathol 2020, 33:2128–2138.

32. Baseler L, de Wit E, Scott DP, Munster VJ, Feldmann H: Syrian hamsters (Mesocricetus auratus) oronasally inoculated with a Nipah virus isolate from Bangladesh or Malaysia develop similar respiratory tract lesions. Vet Pathol 2015, 52:38–45.

33. Prescott J, Safronetz D, Haddock E, Robertson S, Scott D, Feldmann H: The adaptive immune response does not influence hantavirus disease or persistence in the Syrian hamster. Immunology 2013, 140:168–178.

34. Liu J, Li S, Liu J, Liang B, Wang X, Wang H, Li W, Tong Q, Yi J, Zhao L, Xiong L, Guo C, Tian J, Luo J, Yao J, Pang R, Shen H, Peng C, Liu T, Zhang Q, Wu J, Xu L, Lu S, Wang B, Weng Z, Han C, Zhu H, Zhou R, Zhou H, Chen X, Ye P, Zhu B, Wang L, Zhou W, He S, He Y, Jie S, Wei P, Zhang J, Lu Y, Wang W, Zhang L, Li L, Zhou F, Wang J, Dittmer U, Lu M, Hu Y, Yang D, Zheng X: Longitudinal characteristics of lymphocyte responses and cytokine profiles in the peripheral blood of SARS-CoV-2 infected patients. EBioMedicine 2020, 55:102763.

35. Wang F, Nie J, Wang H, Zhao Q, Xiong Y, Deng L, Song S, Ma Z, Mo P, Zhang Y: Characteristics of Peripheral Lymphocyte Subset Alteration in COVID-19 Pneumonia. J Infect Dis 2020, 221:1762–1769.

36. Liu J, Liu Y, Xiang P, Pu L, Xiong H, Li C, Zhang M, Tan J, Xu Y, Song R, Song M, Wang L, Zhang W, Han B, Yang L, Wang X, Zhou G, Zhang T, Li B, Wang Y, Chen Z, Wang X: Neutrophil-to-lymphocyte ratio predicts critical illness patients with 2019 coronavirus disease in the early stage. J Transl Med 2020, 18:206.

37. Tan L, Wang Q, Zhang D, Ding J, Huang Q, Tang Y-Q, Wang Q, Miao H: Lymphopenia predicts disease severity of COVID-19: a descriptive and predictive study. Signal Transduct Target Ther 2020, 5:33.

38. Diao B, Wang C, Tan Y, Chen X, Liu Y, Ning L, Chen L, Li M, Liu Y, Wang G, Yuan Z, Feng Z, Zhang Y, Wu Y, Chen Y: Reduction and Functional Exhaustion of T Cells in Patients With Coronavirus Disease 2019 (COVID-19). Front Immunol 2020, 11:827.

